# Interrogation of genes controlling biofilm formation using CRISPR interference in *Pseudomonas fluorescens*

**DOI:** 10.1101/476366

**Authors:** Marie-Francoise Noirot-Gros, Sara Forrester, Grace Malato, Peter E. Larsen, Philippe Noirot

**Affiliations:** Biosciences Division, Argonne National Laboratory, Lemont, IL, United States.; Department of Bioengineering, University of Illinois at Chicago, Chicago, IL, United States.

## Abstract

Bacterial biofilm formation involves multigenic signaling and regulatory pathways that control the transition from motile to sessile lifestyle, production of extracellular polymeric matrix, and maturation of the biofilm complex 3D structure. Biofilms are extensively studied because of their importance in biomedical, ecological and industrial settings. Genetic approaches based on gene inactivation are powerful for mechanistic studies but often are labor intensive, limiting systematic gene surveys to the most tractable bacterial hosts. Here, we adapted the CRISPR interference (CRISPRi) system for use in *P. fluorescens*. We found that CRISPRi is applicable to three genetically and physiologically diverse species, SBW25, WH6 and Pf0-1 and affords extended periods of time to study complex phenotypes such as cell morphology, motility and biofilm formation. In SBW25, CRISPRi-mediated silencing of the GacA/S two-component system and genes regulated by cylic-di-GMP produced phenotypes similar to those previously described after gene inactivation in various *Pseudomonas*. Combined with detailed confocal microscopy of biofilms, our study also revealed novel phenotypes associated with biofilm architecture and extracellular matrix biosynthesis as well as the potent inhibition of SBW25 biofilm formation mediated by the PFLU1114 protein. Thus, CRISPRi is a reliable and scalable approach to interrogate gene networks in the diverse *P. fluorescens* group.

## Introduction

Biofilms are the prevalent state of bacterial life in nature^1^ and biofilm formation is an integral part of the prokaryotic life cycle. Biofilms are clusters of microorganisms embedded in a self-produced matrix of extracellular biopolymers that provide shelter, allow cooperation between bacterial cells, interactions with the environment, and confer the ability to colonize new niches by dispersal of microorganisms from the microbial clusters ^2,3,4^. Biofilms can form on virtually any abiotic or biotic surface. Often associated with chronic infections and resistance to antibiotic treatments, biofilms are generally considered harmful to human health ^5,6^. In contrast, biofilm-forming non-pathogenic bacteria can effectively protect from infection by pathogens^7^,and can promote the growth of plants and stimulate symbiotic interactions between mycorrhizal fungi and plant roots ^8-12^. Thus, there is considerable interest in deciphering at a molecular level the regulatory mechanisms of biofilm formation and dispersion to combat biofilms but also to better control them.

Biofilm formation is triggered by environmental cues and involves coordinated responses from a number of cellular processes such as flagellar assembly and secretion of extracellular polymeric substances (EPS). In bacteria, two-component systems (TCS) sense environmental stimuli and translate this information into cellular responses through coordinated regulation of genetic programs ^13-15^. In Gram-negative bacteria, the well-characterized GacA/S TCS regulates the expression of genes involved in quorum sensing, stress responses, biofilm formation and virulence ^16,17^. The GacA/S system is composed of a membrane-bound sensor histidine kinase GacS and its cognate response regulator GacA^16^. In γ-proteobacteria such as *Pseudomonas* and *Halomonas*, inactivation by mutation of either GacA or GacS dramatically affects the production of EPS, secondary metabolites and iron homeostasis^18-21^. EPS are mainly composed of alginate and exopolysaccharides and contribute to the biofilm architecture. EPS production is regulated by the Gac/Rsm signaling cascade, involving GacA/S and the non-coding small regulatory RNAs RsmZ and RsmY^17,18,22-24^. In addition, this signaling cascade regulates about 700 genes involved in a wide range of biological functions, including biofilm formation and oxidative stress response^17,24,25^.

Cyclic diguanylate (c-di-GMP), a near universal intracellular signaling messenger, regulates many aspects of bacterial growth and behavior ^26,27^, including the transition from motile-to-sessile lifestyle leading to biofilm formation and virulence^28,29^. Intracellular levels of c-di-GMP can be modulated through the balanced activities of two classes of enzymes: (i) the diguanylate cyclases (DGCs) that synthetize c-di-GMP from two GTPs and

(ii) the diguanylate phosphodiesterases (PDEs) that degrade c-di-GMP in pGpG and GMP ^28,29^. In addition to DCGs (which contain GGDEF domains) and PDEs (which contain EAL or HY-GYP domains), bacterial genomes generally encode multiple c-di-GMP-binding proteins, including PilZ-domain effectors and proteins with ‘degenerate’ GGDEF/EAL domains that are enzymatically inactive but still bind c-di-GMP ^30,31^. Although the propensity of bacteria to form biofilm correlates with higher intracellular concentrations of c-di-GMP^32^, it is not currently understood how a subset of the multiple DGCs and PDEs can act to produce a specific phenotypic outcome^26^. Extensive studies in *P. aeruginosa* uncovered the roles of many DGCs, PDEs and c-di-GMP-binding effectors during biofilm development, from the initial stage of cell adhesion on a surface to colony formation and biofilm maturation and dispersion^28^. In *P. aeruginosa*, the initial biofilm formation stage is governed by the di<u>g</u>uanylate <u>c</u>yclase promoting <u>b</u>iofilm GcbA that regulates surface attachment via modulation of the flagellum-driven motility^33^. GcbA is also involved in biofilm dispersion through post-translational processing of the chemosensory protein BdlA, known to regulate PDE activities such as that of the DipA protein ^34-36^. The PDE BifA modulates biofilm formation and motility by altering exopolysaccharide production and flagellar function ^37-39^. C-di-GMP also regulates alginate synthesis through binding to the PilZ-type domain of alginate co-polymerase Alg44. Additional levels of regulation involve the PDE protein RimA which modulates the activity of RimK, an enzyme that modifies the ribosomal protein RpsF by adding glutamate residues to its C terminus, thus altering ribosome abundance and function ^40,41^. Altogether, these studies reveal that c-di-GMP-binding enzymes and proteins act in a coordinated manner to control the various stages of the planktonic-to-biofilm transition, altering motility, promoting cell adhesion, producing EPS and shaping biofilm architecture.

Here, we focus on studying c-di-GMP-associated regulators that control biofilm formation in the rhizobacterium *P. fluorescens*. Soil and plant-associated bacteria such as *P. fluorescens* isolates can provide beneficial ecological services to various plants ^42-44^ and form dynamic and highly structured biofilm at aspen roots^11^. Genomic comparisons between multiple *P. fluorescens* strains revealed a high genomic heterogeneity with a conserved core genome and a widely diverse pan-genome encoding specialized activities, many being relevant for colonization of the rhizosphere and plant-bacteria interactions^45,46^. *P. fluorescens* genomes typically encode about 50 proteins with c-di-GMP-binding signatures, including homologs for GbcA, BifA, DipA and Alg44. Extensive phenotypic screens have been performed in *P. fluorescens* to identify genes controlling biofilm formation. In *P. fluorescens* SBW25, screens for wrinkly spreader biofilm phenotypes, which are associated with enhanced formation of cellulose-based matrix at the air-liquid (AL) interface, identified many DCGs and associated regulators involved in c-di-GMP homeostasis^47-51^. In *P. fluorescens* Pf0-1, a systematic survey of knockout mutants revealed that about a third of c-di-GMP-associated proteins exhibit strong biofilm phenotypes across many growth conditions whereas the remaining mutants exhibited weak phenotypes in only a small number of conditions ^52^. However, the biological roles of most the c-di-GMP-binding proteins still remain to be characterized. Understanding these biological roles relies on our ability to associate specific c-di-GMP-related proteins with particular biofilm phenotypes. Most previous studies, however, have been based on a colorimetric assay that measures EPS accumulation in mature biofilms ^53^. While this assay is rapid and reliable, it cannot report on the full range of phenotypes (e.g., cells abundance, EPS architecture) that characterize biofilm development.

In this work, we used a CRISPRi-based approach to investigate the role of genes belonging to the c-di-GMP regulatory network in biofilm formation and architecture. CRISPRi can be used to modulate gene expression and to study genes essential for cell survival ^54,55,56^. Here, we adapted the CRISPRi system to *P. fluorescens* SBW25 and validated its application for gene silencing in SBW25, WH6 and Pf0-1 strains. In SBW25, we show that CRISPRi allows to study phenotypes quantitatively at the cell, colony and biofilm levels. Then, we applied our CRISPRi system to interrogate genes involved in c-di-GMP signaling pathways that control biofilm formation and cell motility. We report that CRISPRi-mediated gene silencing is a robust and reliable approach for quantitative phenotyping of complex bacterial traits such as swarming motility and biofilm mass, structure and composition that can be used to discover the function of uncharacterized genes.

## Results

### CRISPRi mediates efficient gene silencing in *P. fluorescens*

In the CRISPRi system, a small guide RNA (gRNA) directs the catalytically inactive dCas9 protein to bind at or near a promoter region and sterically hinder the initiation or elongation of transcription, resulting in silencing of gene expression ^54,55^. CRISPRi systems have been previously shown to sterically block transcription of genes in the model bacteria *E. coli* and *B. subtilis* ^55,56^. We adapted the CRISPRi system for *P. fluorescens* by constructing a system comprised of two compatible plasmids (Supplementary Fig. S1). One plasmid carries the *S. pyogenes* dCas9 gene under control of the P*TetA* promoter that can be induced by the presence of anhydrotetracyclin (aT)^57^ in the growth medium. The other plasmid constitutively expresses a gRNA (see Materials and Methods).

We assessed the functionality of our CRISPRi/dCas9 system in three different *P. fluorescens* strains: SBW25, WH6 and Pf0-1 that expressed a chromosomal copy of the *mNG* gene, encoding the mNeonGreen fluorescent protein under control of a constitutive Pc promoter^11^. This construct was inserted at similar chromosomal locations in all strains^11^. We designed two pairs of gRNAs, one pair (Pc4 and Pc5) targeting transcription initiation at the Pc promoter and the other pair (Pc2 and Pc3), targeting transcription elongation at a site overlapping the start of the open reading frame (ORF, Supplementary Fig S2, Supplementary Table S1). These RNA guides target DNA sites either copying the template (T, Pc3 and Pc4) or non-template (NT, Pc2 and Pc5) strand (Supplementary Fig. S2). Upon induction of dCas9 expression, the effect of each gRNA on fluorescence intensity was monitored over time using flow cytometry (Figure 1, Supplementary Fig. S2). In this survey, we found that gRNAs Pc4 and Pc5 targeting the transcription initiation of *mNG* gene resulted in the highest decrease of fluorescence relative to the control without gRNA. However, part of this decrease is observed in the absence of inducer (T=0, Supplementary Figure. S2), suggesting that dCas9 is expressed at some basal level. The basal expression of dCas9 was found to have minimal effects with the Pc2 gRNA targeting transcription elongation and copying the NT strand (gRNA_NT_) in SBW25 and WH6 but not in Pf0-1 (Supplementary Figure. S2).

**Figure 1.**
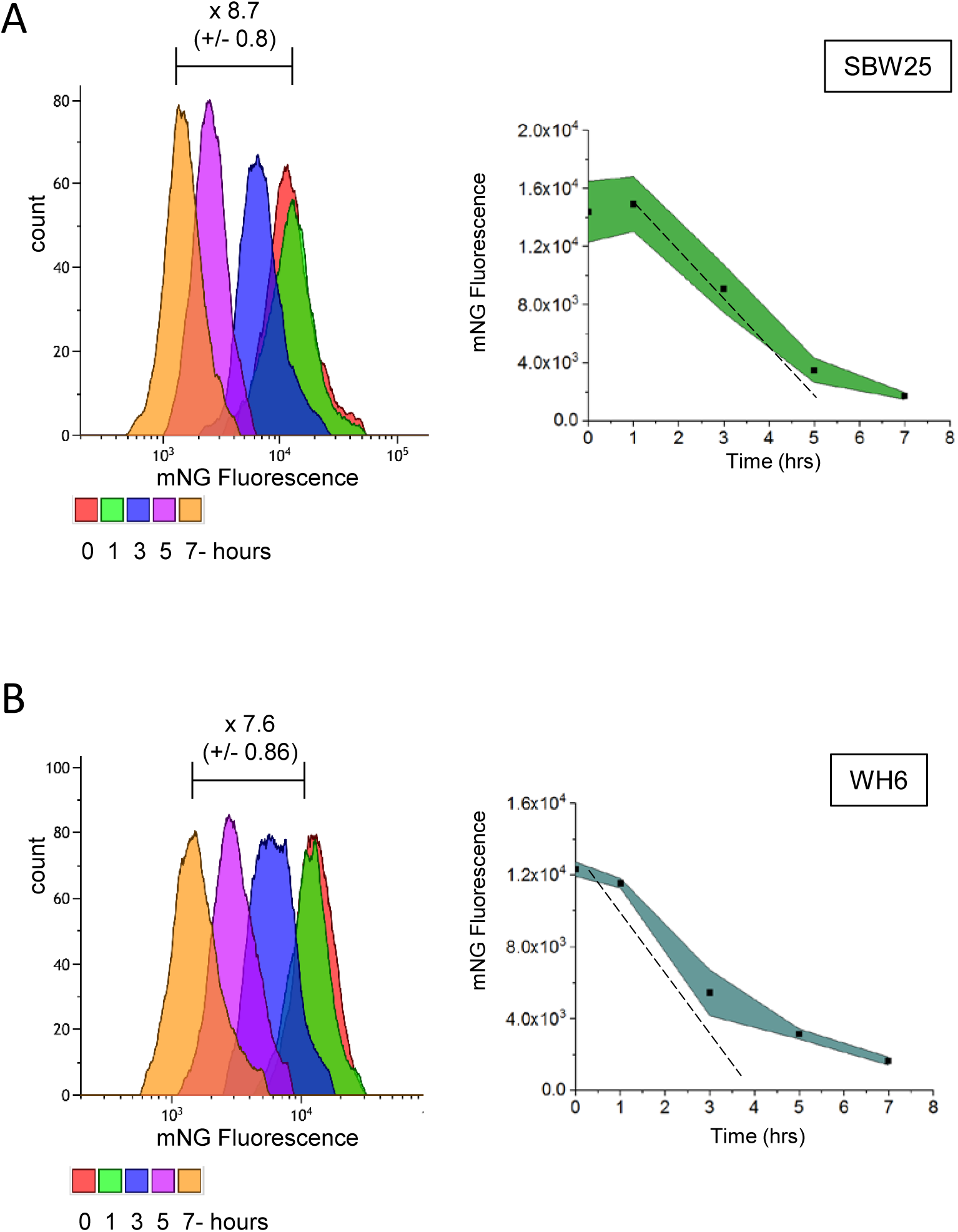
Gene silencing by dCas9-gRNA system in *P. fluorescens*. SBW25 (A) and WH6 (B) strains contain an identical DNA cassette coding for the mNG fluorescent protein expressed from a constitutive promoter (Pc) inserted at a similar chromosomal locus. Both strains harbor the pPFL-Cas9 plasmid and the pPFL-gRNA plasmid expressing the gRNA Pc2 targeting transcription elongation of the mNG gene at a site overlapping the start of the ORF (see also Fig. S2). Green fluorescence intensities were monitored over 7 hours after induction of dCas9 (illustrated by colors, right panels). Upon silencing, fluorescence decreases overtime due to dilution from cell divisions (left panels). Dotted lines represent the expected dilution of the GFP in SBW25 (doubling time 135 min) and WH6 (doubling time 120 min).

Based on these findings, we chose to design our next experiments using (i) gRNANT guides that target the start of the ORFs of our genes of interest and (ii) a strain harboring the two plasmids and expressing dCas9 but no gRNA as a control for no CRISPRi activity. We evaluated the kinetic of fluorescence decrease caused by elongation-blocking gRNANT in SBW25 and WH6 (Figure 1). A complete block of *mNG* expression is expected to result in a 50% decrease of the fluorescence intensity after each cell doubling, owing to the short maturation time and high stability of the mNG protein^58^. In the conditions of our assay, the generation times for SBW25 and WH6 were 135 min and 120 min, respectively. For both strains, the observed decrease of fluorescence is only slightly slower than the expected dilution of the mNG protein from cell division, indicating a strong although incomplete silencing (Figure 1). Monitoring of the effect of targeting the T strand for elongation transcription block by Pc3 gRNA revealed a lower decrease of fluorescence (3.2-fold after 7 hours) compared to the non-induced condition (Supplementary Fig. S2, Supplementary Table S2). This result corroborates previous studies highlighting the importance of using the NT strand for optimal repression efficiency ^54,55^. However, the T strand could still be used to provide intermediate levels of gene down regulation.

### CRISPRi-mediated silencing of essential genes involved in cytokinesis and morphogenesis

To assess the efficacy of our CRISPRi system to generate observable phenotypes in *P. fluorescens*, we interrogated the *ftsZ* and *mreB* genes that are essential for bacterial survival and were previously shown to produce characteristic defects in cell division or cell shape upon mutation. Upon depletion of tubulin-like protein FtsZ, bacterial cells typically grow as long and non-septate filaments resulting from failure to assemble a functional FtsZ division ring ^59-62^. The depletion of actin-like MreB in *E. coli* cells results in loss of the rod-shaped morphology and gives rise to enlarged round cells ^63^. When there are several MreB-like proteins, such as in *B. subtilis*, the depletion of each homolog results in aberrant cell morphology phenotypes such as wider, inflated and twisted cells, ultimately leading to cell lysis ^64,65^.

*P. fluorescens* SBW25 *ftsZ* (PFLU0952) and *mreB* (PFLU0863) genes, which are predicted to be monocistronic (Supplementary Fig. S3), were subjected to CRISPRi-mediated silencing by expressing a gRNA targeting *ftsZ* (*ftsZ_NT_*) or mreB (*mreB_NT_*) in cells induced for dCas9 expression. After 5 hours of incubation at 25°C in the presence of inducer (aT 0.1μg/ml), cells expressing the *ftsZ_NT_* guide exhibited a characteristic cell filamentation phenotype (Figure 2) consistent with *ftsZ* knockdown. The morphological defects appeared 3 hours after induction, involved all observed cells after 5 hours, and this phenotype persisted in overnight cultures (18 hours) (Supplementary Fig. S4). Cells expressing the *mreB_NT_* guide exhibited the characteristic morphological defects associated with a defect in *mreB* expression, including inflated and round cells with part of them bursting in overnight cultures (Figure 2, Supplementary Fig. S4). Of note, we found that the expected morphological defects could also be observed using gRNAs copying the T-strand and targeting transcription elongation sites of *ftsZ* and *mreB* genes, under conditions where inducer (aT) concentration is 5-fold higher and incubation times are longer (Supplementary Fig. S5). These findings confirm the importance of using the non-template strand for gRNAs displaying optimal gene repression. They also indicate that the template strand can be used when a milder repression is needed. Altogether, our results show that CRISPRi is suitable for gene silencing associated with phenotypic analyses over several hours in *P. fluorescens*.

**Figure 2.**
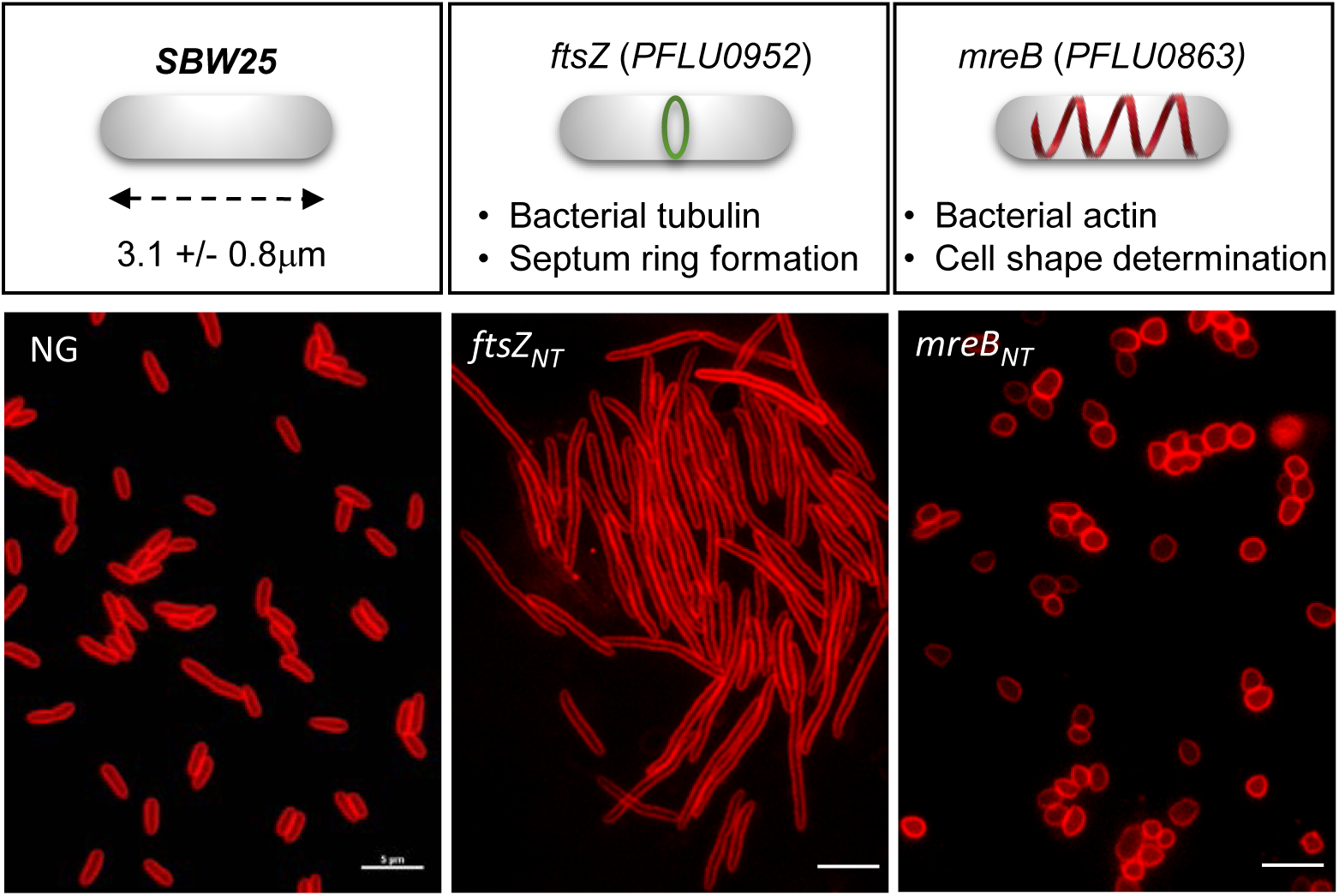
CRISPRi-mediated silencing of genes involved in cell division and cell morphology in *P. fluorescens*. Cells harboring the pPFL-Cas9 plasmid and the pPFL-gRNA plasmid expressing a gRNA that copies the non-template strand at the start of the ORFs to block transcription elongation. (A) Control with no guide, (B) Silencing of *ftsZ* (PFLU0952) using a ftsZ_NT_ guide, and (C) Silencing of *mreB* (PFLU0863) using a mreB_NT_ guide. Strains were grown 5 hours in the presence or absence of inducer (aT 0.1 μg/ml). Cells were stained with the membrane fluorescent dye FM-4-64 prior to observation by epifluorescence microscopy. Scale bars indicate 2μm.

### Silencing of the two-component sensor kinase GacS impairs mobility, biofilm formation and improves tolerance to acute oxidative stress

CRISPRi system has been shown to confer a rapid gene silencing that is stable over time ^54,66^. As numerous *P. fluorescens* phenotypes related to mobility and biofilm are typically measured after 48 hours, we investigated CRISPRi-mediated silencing of the pleiotropic *gacS* (PFLU3777) gene. We found that upon induction of *dCas9* expression, the strain expressing the *gacS_NT_* guide was totally impaired in swarming motility after 48 hours (Figure 3), in keeping with the tight control of motility by *gacS* in *Pseudomonas* ^17,24,25,67,68^. Expression of the *gacS_T_* guide also led to a clear defect in swarming (supplementary Fig. S6). Silencing of *gacS* also produced strong defects in biofilm formation relative to the control strain with no gRNA (Figures 4-6). The biofilm pellicles formed at the air-liquid interface were imaged by confocal microscopy after staining of the cells with the membrane-specific dye FM1-43 biofilm tracer (green) or staining of the exopolysaccharides with the Congo red dye (red). A 3D reconstruction of the biofilm revealed that *gacS* silencing produced a thinner and less cohesive pellicle with lower cell biomass compared to control (Figure 4, 5). Likewise, the exopolysaccharide matrix exhibited irregular density and discontinuities (Figure 4, 5), indicating impaired biofilm formation and altered physical properties of the pellicle. Thus, CRISPRi-mediated silencing of *gacS* fully reproduced the phenotypic defects in surface motility and biofilm formation that are hallmarks of *gacS* mutants in *Pseudomonas^17,18,20,24,25^*. Interestingly, the *gacS* knockdown strain appeared about 3-fold more tolerant to H_2_O_2_ exposure than the control strain (Supplementary Fig. S7). This is in contrast with previous observations in SBW25 that growth of a *gacS*::Tn5 mutant is inhibited in the presence of H_2_O_2_ 25 and may be due to differences in H_2_O_2_ treatments. The increased survival to oxidative stress observed here could result from a short and acute exposure to H2O2 which differs from the long and chronic exposure condition used in previous studies ^25^. From these results, we conclude that CRISPRi-mediated gene silencing enables the reliable study of plate-based phenotypes that take 48 hours to develop. It is therefore a robust approach to interrogate gene-phenotype relationships in *P. fluorescens*.

**Figure 3.**
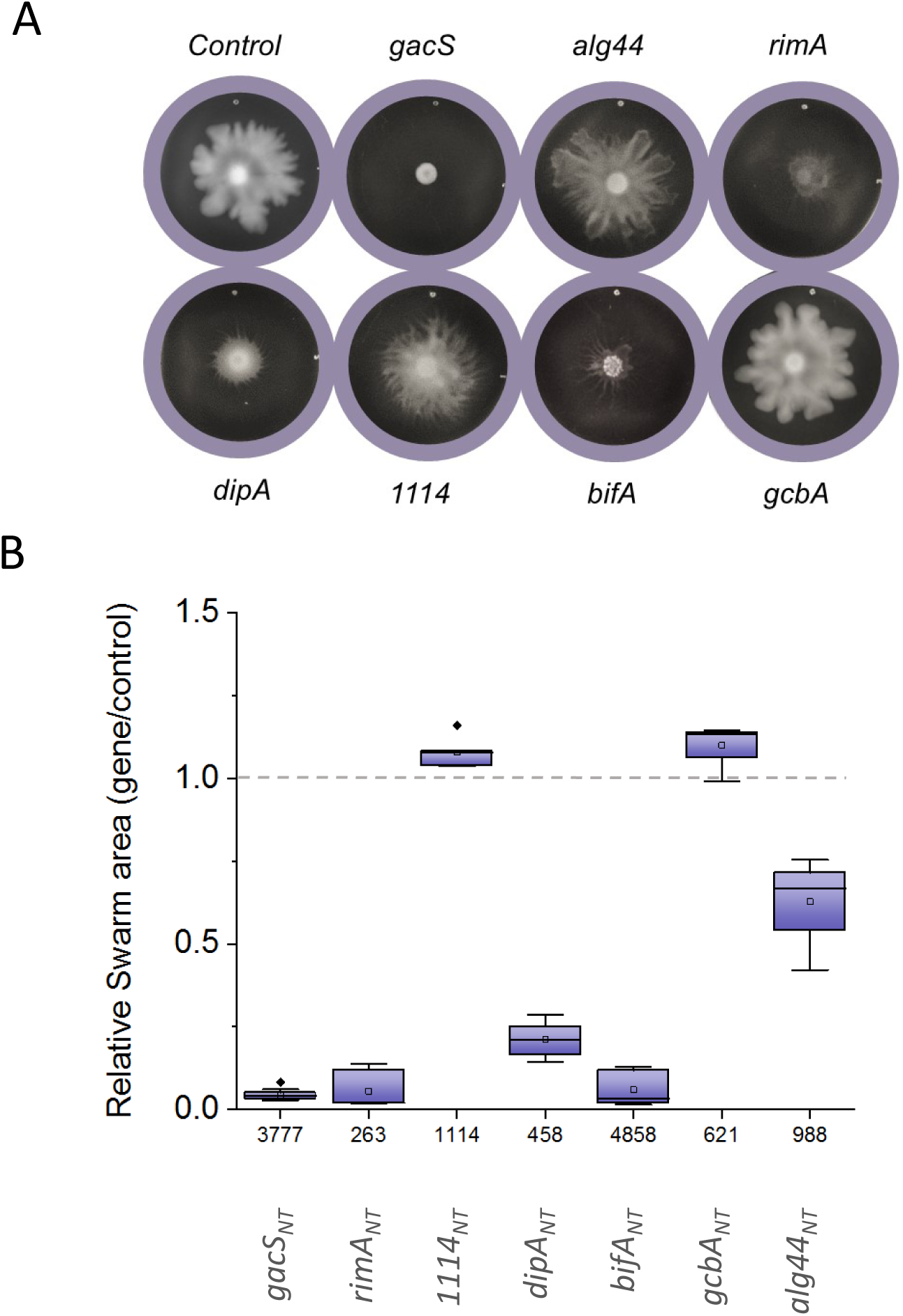
Phenotyping of genes involved in swarming motility. *P. fluorescens* strains harboring plasmids pPFL-Cas9 and pPFL-gRNA, the latter expressing a gRNA that copies the non-template strand (NT) at the start of the ORF to block transcription elongation. Targeted genes include the *gacS* (PFLU3777) encoding the kinase sensor protein GacS and genes encoding c-di-GMP binding proteins RimA (PFLU0263, PDE), DipA (PFLU0458, PDE), the alginate co-polymerase Alg44 (PFLU0988), GcbA (PFL0621, DGC) and BifA (PFLU4858, PDE) (see also Fig. S6). The control corresponds to the pPFL-gRNA plasmid with no guide RNA inserted. (A) Typical swarming morphotypes observed at the surface of soft-agar plates incubated 48 hrs at 25ºC. (B) Box plot representing the swarm areas of cells with silenced gene relative to the control strain (n≥4).

**Figure 4.**
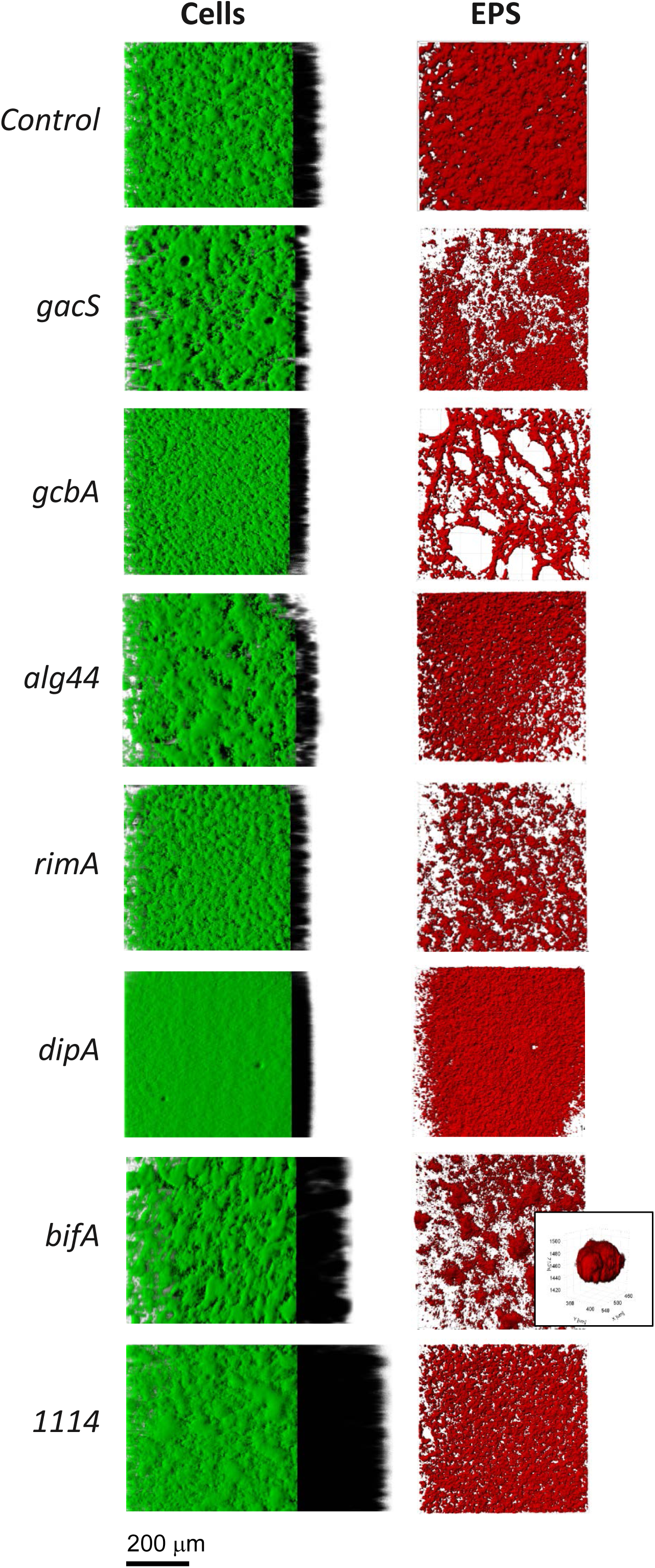
3D-projections of biofilm structures and EPS matrix. *P. fluorescens* strains harboring plasmids pPFL-Cas9 and pPFL-gRNA, the latter expressing a gRNA_NT_ targeting genes *gacS* (PFLU3777, two-component sensor kinase), *rimA* (PFLU0263, PDE), *dipA* (PFLU0458, PDE), *alg44* (PFLU0988, alginase co-polymerase Alg44), *gcbA* (PFL0621, DGC) and *bifA* (PFLU4858, PDE). The control corresponds to the pPFL-gRNA plasmid with no guide RNA inserted. A) 3D architectures of biofilms featured in Fig. 4 including virtual shadow projections of thickness on the right. B) Biofilms grown in the presence of the amyloid dye Congo Red to reveal the structure of EPS.

**Figures 5.**
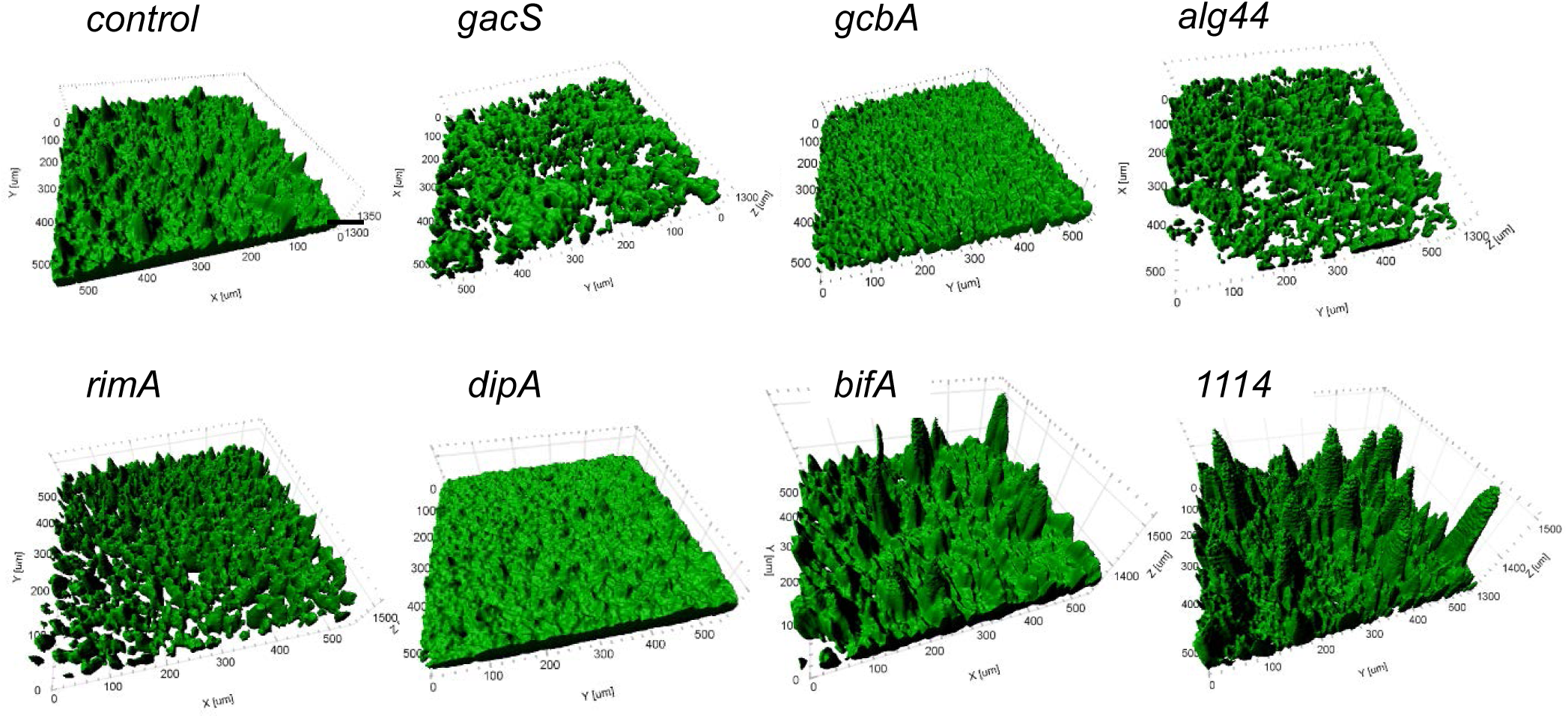
Analysis of biofilm phenotypes upon down regulation of genes involved in GacS and c-di-GMP signaling. Surface rendering (IMARIS software) of reconstituted 3D confocal volume images of AL biofilm pellicles displayed in Figure 4. Targeted genes include the *gacS* (PFLU3777) encoding the kinase sensor protein GacS and genes encoding c-di-GMP binding proteins *rimA* (PFLU0263, PDE), *dipA* (PFLU0458, PDE), *alg44* (PFLU0988) encoding the alginase co-polymerase Alg44, *gcbA* (PFL0621, DGC) and *bifA* (PFLU4858, PDE). The control corresponds to the pPFL-gRNA plasmid with no guide RNA inserted.

### CRISPRi interrogation of genes involved in c-di-GMP signaling for swarming motility and biofilm formation

Next, we applied CRISPRi silencing to five biofilm-associated genes in SBW25 encoding c-di-GMP binding proteins that are known to act at various stages of the biofilm formation in *Pseudomonas^28^*. PFLU0988 is a homolog of *P. aeruginosa alg44* encoding an alginate co-polymerase ^69^. PFLU0621 is a homolog to *P. aeruginosa gcbA* encoding the DGC-promoting biofilm enzyme GcbA^33^. PFLU0458 and PFLU4858 are homologs of *dipA* and *bifA,* respectively, which encode enzymes with characterized PDE activity in *P. aeruginosa and P. putida* ^35,37-39^. PFLU0263 encodes the single EAL domain protein RimA with characterized PDE activity in SBW25 ^41^. In addition, we investigated PFLU1114, a gene of unknown function, encoding a putative PDE (Pseudomonas Ortholog Group POG020457). Of note, *rimA* is the first gene of an operon comprising *rimB* (PFLU0262) and *rimK* (PFLU0261), suggesting that CRISPRi-mediated silencing of *rimA* will likely also affect expression of the downstream genes (Supplementary Fig. S3). For each selected gene, we designed a gRNA_NT_ targeting a site at the start of the ORF and tested the effect of gene silencing on swarming motility and biofilm formation.

Silencing of *rimA*, *bifA* and *dipA* drastically reduced swarming ability whereas silencing of *gcbA*, *alg44* and PFLU1114 did not substantially affect swarming (Figure 3). The swarming phenotypes obtained upon silencing with gRNA_T_ were consistent but of reduced amplitude relative to those obtained with gRNA_NT_ (Supplementary Fig. S6). To assess biofilm formation, bacterial cells and EPS present in air-liquid pellicles were dyed separately and imaged by confocal microscopy (Figure 4). Reconstructed 3D surface renderings (Figure 5) were analyzed for thickness, roughness and substratum coverage, indicative of biomass, structure and fraction of the substratum covered, respectively (Figure 6). Biofilms formed by RimABK-depleted cells were thinner, flakier and less rough than the control with an irregular EPS matrix structure (Figure 4). These observations are in agreement with the loss of biofilm ability previously observed in a SBW25 ΔPFLU0263 strain^41^. Biofilms formed by DipA-depleted cells appear very flat and dense with a high substratum coverage as well as a homogenous repartition of EPS (Figures 4-6). In contrast, the depletion of BifA gave rise to a substantial increase of biofilm cell mass, thickness, roughness and formation of clumps in the EPS matrix (Figures 4-6). The largest increase in biofilm cell mass, thickness and roughness was observed upon depletion of PFLU1114, which did not alter the texture of the EPS matrix (Figure 4) but produced particularly thick and rough biofilms (Figure 5). Depletion of the alginate co-polymerase Alg44 resulted in a thinner biofilm with more dispersed EPS matrix than the control (Figure 4), in keeping with its role in synthesis of the exopolysaccharide alginate^70^. Finally, the depletion of GcbA slightly impaired biofilm cell mass, thickness and architecture but drastically altered the EPS matrix production and cohesion (Figures 4-6). GcbA has previously been described as a regulator of the initial step of cell adhesion during biofilm formation^33^. Our findings suggest an additional role of GcbA in the regulation of EPS matrix formation in SBW25.

**Figure 6.**
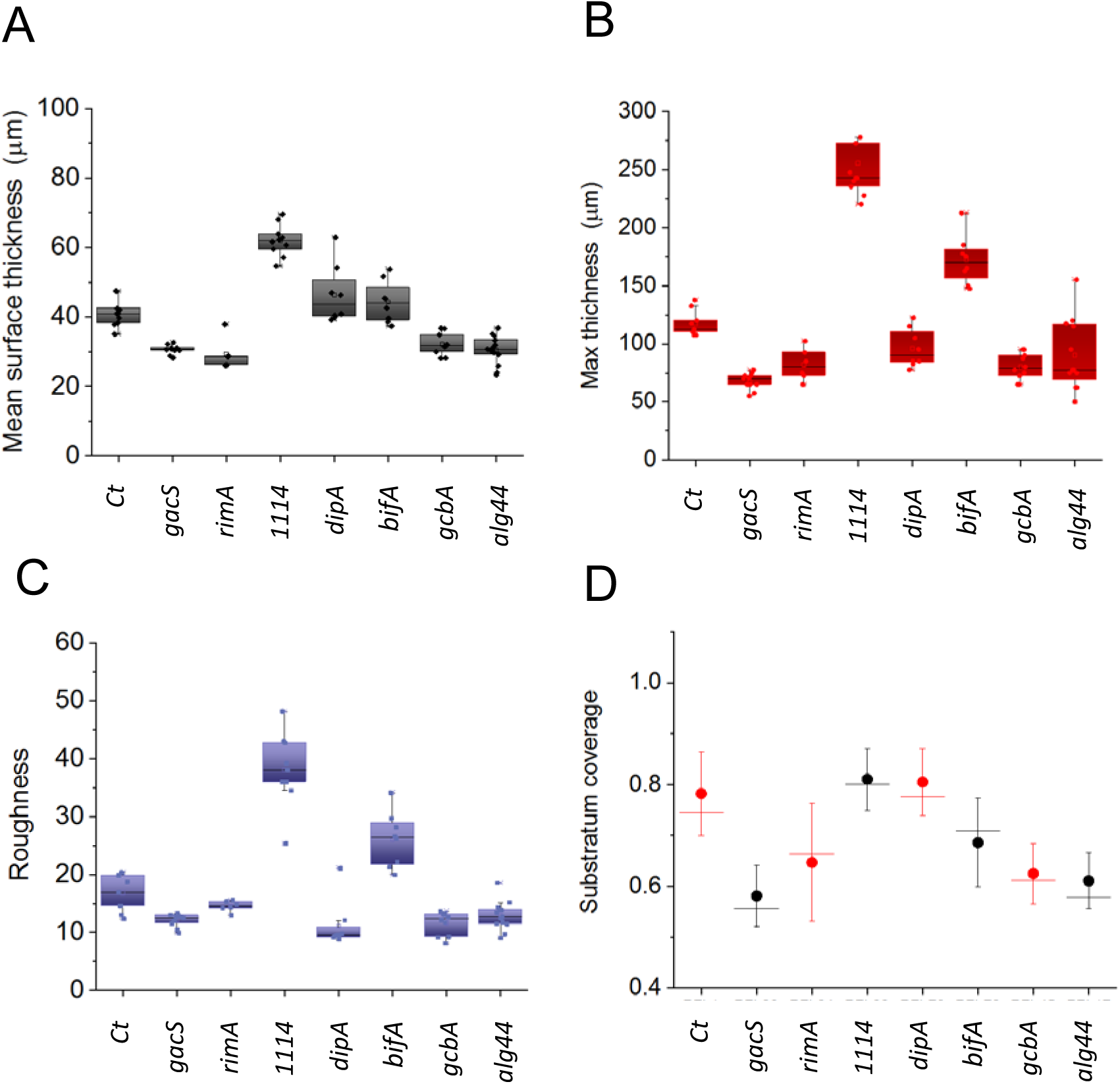
Biofilms parameters upon down regulation of genes of c-di-GMP pathway. Quantification of the 3D reconstructed AL pellicles biofilm volumes (see Fig. 5) were performed using IMARIS x64 9.0.2 XTension software package. Box plots display the distribution of data obtained from observation of a biofilm pellicle at different places (n≥6) for. A) Mean thickness biovolumes (μm); B) Maximum thickness biovolumes (μm); C) Roughness coefficient of biofilms and D) Biofilm surface coverage ratio (1=no gap in surface).

## Discussion

We adapted the CRISPRi system to *P. fluorescens* in three genetically and physiologically diverse *P. fluorescens* species: SBW25, WH6 and Pf0-1^71^. We found that CRISPRi effectively inhibited the expression of a constitutively expressed fluorescent reporter in all three species, suggesting that our system can be utilized across the *P. fluorescens* group. The amplitude of the inhibition depended on the DNA site and DNA strand targeted by the gRNA (Supplementary Fig. S2) and the concentration of aT inducer (Supplementary Fig. S5), demonstrating that these features can be used to modulate the level of inhibition. We elected to design gRNAs that bind to the start of the ORFs and are complementary to the protein coding strand because in our assay these features yielded i) a very low leaky repression in the absence of inducer (in SBW25 and WH6), ii) a substantial inhibition of reporter expression in the presence of inducer and importantly, iii) a homogeneous distribution of fluorescence intensity in the cell population (Figure 1). These findings indicate that the induction of *dCas9* blocks transcription elongation resulting in efficient gene silencing.

The silencing capacity of CRISPRi was further investigated in SBW25 using a variety of genes linked to specific phenotypes including cell division and cell morphology that are observed after 3-18 hours of growth, as well as complex phenotypes associated with the regulation of bacterial motility and biofilm formation that are scored after 48 hours on plate and liquid culture assays. We found that CRISPRi-mediated silencing of *ftsZ* and *mreB* genes caused characteristic cell morphology defects in the whole cell population and silencing of genes involved in signaling pathways that control swarming motility and biofilm formation produced phenotypes that are consistent with previously observed phenotypes in gene inactivation mutants. Our analysis also revealed novel phenotypes related to the EPS matrix for previously studied genes (e.g., *gcbA*, *bifA*), and allowed the discovery of a key role of PFLU1114 in inhibiting biofilm formation in SBW25. Recently, a CRISPRi system based on *S. pasteurianus* dCas9 was successfully tested in *P. aeruginosa* and in other *Pseudomonas* but the described leakiness of the promoter controlling *SpadCas9* expression represented a limitation for phenotype analysis in *P. fluorescens ^72^*. Here, our CRISPRi system is a robust and reliable approach to study complex phenotypes related to *P. fluorescens* life styles and behaviors, potentially opening the way to more systematic studies in this diverse bacterial group.

In *P. fluorescens* SBW25, we found a strong effect of GacS depletion on swarming, in agreement with previous observations that swarming motility was reduced in a *gacS::Tn5* mutant ^25^. In *Pseudomonas* and other γ-proteobacteria, the sensor kinase GacS is at the heart of the regulation of genes involved in biofilm formation and defects in swarming and biofilm formation are the hallmark of *gacS* mutants. In response to environmental stimuli, GacS phosphorylates and activates of the response regulator GacA leading to the upregulation of the expression of regulatory small RNAs. These sRNAs bind to the global regulator RsmA, releasing its repression of genes necessary for biofilm formation^20,40^. The decrease in biofilm formation observed upon CRISPRi-mediated silencing of *gacS* is in agreement with the regulatory model and with phenotypes observed in *gacS* mutants in various *Pseudomonas* ^17,18,25,40,68^. In SBW25, GacS is also regulating genes involved in oxidative stress response, and the *gacS::Tn5* mutant has a reduced capacity to survive exposure to sub-lethal doses of H_2_O_2_^25^. This greater sensitivity to H_2_O_2_ was correlated with a reduction in the expression of *katE* encoding a catalase^25^.

Upon silencing of *gacS* in our assay, we did not observe such an increased sensitivity to H_2_O_2_ but rather a slight and reproducible increase in H_2_O_2_ tolerance. Interestingly, a strong upregulation (300-fold) of the superoxide dismutase SodA was also observed in the *gacS::Tn5* mutant ^25^, suggesting that the higher tolerance we observe upon GacS depletion may result from the short and acute exposure to H_2_O_2_ we applied versus the chronic, low-dose exposure to H_2_O_2_ applied in the previous study^25^.

Our analysis of genes from the c-di-GMP regulatory network, involved in different stages of biofilm formation, revealed that CRISPRi-mediated silencing can be used for fine dissection of biofilm phenotypes and gene function study. We found that the down regulation of the DGC *gcbA* affects biofilms thickness and structure as well as ESP production. Targeting of the *rim* operon strongly impaired swarming motility, a phenotype also found associated with a deletion of *rimK* in various *Pseudomonas*^41^. In SBW25, swarming is slightly affected^41^ by *ΔrimK* deletion while not affected at all by *ΔrimA* and *ΔrimB* deletions^41^. The swarming deficiency we observed upon targeting *rimA* suggests an efficient silencing of the whole the *rimABK* operon. The *rimABK* operon is highly upregulated during the early stages of colonization of the plant roots, and the *ΔrimK* deletion significantly impedes rhizosphere colonization by *P. fluorescens* SBW25^40,41^. Impaired biofilm formation upon silencing of the *rim* operon observed here could reflect a defect in the ability to colonize surfaces.

In *Pseudomonas*, GcbA plays a key role in transition to irreversible attachment to surface, a process linked to EPS production^33^. CRISPRi-mediated silencing of *gcbA* also resulted in a decrease of biofilm mass and thickness of the biofilm, in keeping with the reduction of biofilm biomass observed in *P. fluorescens* Pf0-1 *ΔgcbA* mutant^73^ We also observed a mild defect in the structure of EPS in the biofilm, consistent with a defect in cell attachment^28,33^. Major changes in biofilm architecture were also observed after depletion of PDEs. The silencing of *bifA* and *dipA* produced extreme but opposite phenotypes related to biofilm mass, thickness and structure. These observations confirmed that biofilm formation is not directly promoted by a higher concentration of c-di-GMP in the cell but is regulated by discrete and interconnected pathways that respond to local concentrations of c-di-GMP ^26^. The DipA protein plays a crucial role in biofilm formation and dispersion in *P. aeruginosa* as a *ΔdipA* mutant exhibits reduced swarming motility, increased EPS production and reduced cell dispersal^35^. Our observations are in line with these results and further reveal the flatness and increased density of biofilms formed by DipA-depleted cells, potentially accounting for the reduced ability for cell dispersal. In contrast, we found that the depletion of BifA leads to a large increase in biofilm thickness and 3-dimensional architecture with the formation of mushroom cap-shaped clumps in the ESP matrix. This corroborates the phenotypes of a *ΔbifA* mutant in *P. aeruginosa*, which exhibits a hyper biofilm phenotype and a loss of swarming ability^38,74^. Our findings also reveal a role of BifA in controlling spatial organization and structure of EPS in SBW25. Finally, we discovered that the depletion of PFLU1114 triggers the formation of biofilms that are remarkably thick and highly structured. Because the swarming motility and the synthesis of EPS were not affected by PFLU1114 depletion, we hypothesize that PFLU1114 acts after the attachment stage to limit the cell density in the maturing biofilm.

We propose that CRISPRi silencing is an appealing approach for future systematic interrogation of gene networks in *P. fluorescens*. Such approaches have been successfully applied in other bacteria to identify essential genes^56,75^. Our CRISPRi system could be applied to investigate systematically all the genes involved in signaling pathways that regulate bacterial life styles, including biofilm formation. A single strain can be transformed with a combination of two plasmids, producing strain derivatives with identical genetic backgrounds. These cells can be propagated without inducer, potentially limiting the selective pressure caused by CRISPRi activity and reducing the probability for spontaneous accumulation of adaptive mutations that restore cell fitness, as observed in cells carrying gene deletions ^76-78^. Upon induction of dCas9, gene silencing takes effect within 1-2 hours and persists over 2 days, enabling to monitor complex phenotypes. This study illustrates the relevance of CRISPRi as a tool to investigate molecular mechanisms and regulatory pathways involved in various environmental bacteria.

## Methods

### Microbial strains and media

*P. fluorescens* strains SBW25, and WH6 and Pf0-1 genetically labelled by the mNeongreen (mNG) fluorescent protein expressed from a constitutive promoter (Pc) were described in a previous study^11^. Plasmids were constructed and propagated in *E. coli* DH5α (Biolabs) prior to transformation in *Pseudomonas*. Bacterial cultures and swarming motility assays were performed in LB medium (liquid or agar-containing), Biofilm assays were performed and in M9 medium supplemented with glucose 0.4% as carbon source. When appropriate, kanamycin (50 μg/ml) and gentamycin (10 μg/ml) were added to select for plasmid maintenance. Anhydrotetracycline (aT) was used as inducer of the P_tetA_/TetR promoter/repressor system at the indicated concentrations. Cells were grown with shaking at 37°C for *E. coli* and 28°C for *P. fluorescens*. Swarming and biofilm assays with *P. fluorescens* were performed in a humidity-controlled growth chamber at 25 °C under 70% humidity.

### Construction of CRISPRi vectors

pPFL-dCas9 was constructed by insertion of a PCR-amplified DNA fragment carrying the minimal replicon sequence of the *P. fluorescens* plasmid pVS1 ^79^ into the BsrGI/StuI restriction sites of plasmid pAN-PTet-dCas9 vector ^57^. The resulting plasmid pPFL-dCas9 can be stably propagated in *P. fluorescens* and maintained by applying kanamycin selection. In pPFL-dCas9, the gene encoding dCas9 is placed under a Tetracyclin-inducible P_tetA_ promoter controlled by the *tetR* repressor gene (Fig. S1). The pPFL-gRNA was built by insertion of a PCR-amplified DNA fragment carrying the constitutive J23119 promoter, dCas9 handle and *S. pyogenes* terminator cassette from plasmid pgRNA-bacteria ^55,80^ into the EcoRI/PpuMI restriction site of plasmid pSEVA643 ^81^. The resulting pPFL-sgRNA plasmid is compatible with pPFL-Cas9 in *P. fluorescens*, can be maintained with gentamycin selection, and serves as backbone for the synthesis of gRNAs (Fig. S1). The gRNA sequences were copied either from the template or to the non-template DNA strands of the targeted genes (Fig. S2). The ‘CasFinder’ software package (https://omictools.com/casfinder-tool) was used to design gRNAs with minimal potential for off-target effects ^82^. The gRNA sequences (Table S1) were synthetized as DNA gBlocks (Integrated DNA technologies, https://www.idtdna.com) and cloned into the EcoRI/SpeI restriction sites of pPFL-gRNA. Plasmid constructs were transformed in *E. coli,* purified and transferred in *P. fluorescens* strains SBW25, WH6 and Pf0-1, using standard electroporation techniques^83^.

### Analysis of CRISPRi-mediated phenotypes

*Fluorescence signal by flow cytometry. P. fluorescens* strains expressing the mNeonGreen fluorescent protein and carrying plasmids pPFL-dCa9 and pPFL-gRNA derivatives were grown at 28°C in LB medium supplemented with kanamycin and gentamycin. Overnight cultures were diluted to OD_600_ 0.1 in fresh LB medium supplemented with the same antibiotics and with or without the inducer anhydrotetracyclin (aT) 100 ng/ml and cultures were grown at 28°C under agitation for 7 hours. Culture samples (10 μl) were taken at 0, 1, 3, 5 and 7 hours, diluted in PBS and subjected to flow cytometry (CytoFlex S, Beckman) to quantify green fluorescence. At least 10^4^ particles were counted for each sample. Computerized gating in forward scatter (FSC) and side scatter (SSC) was used to eliminate cell debris. Histograms were generated using the Kaluza 2.0 software (https://www.beckman.com/coulter-flow-cytometers/software/kaluza).

*Morphological defects by epifluorescence microscopy.* Cells carrying pPFL-dCas9 and pPFL-gRNA derivatives were grown at 28 °C in LB supplemented with kanamycin and gentamycin. Overnight cultures were diluted in fresh LB supplemented with the same antibiotics to OD_600_ 0.1, in the presence or absence of anhydrotetracycline (aT) at final concentrations ranging from 100 - 500 ng/ml. Culture samples were taken at various times and mixed with the membrane dye FM4-64 (Invitrogen) prior to immobilization on glass slides padded with agarose 1.3%. Images were acquired using a fluorescence microscope (Nikon Eclipse Ti) equipped with a Plan Apo λ 100x/1,45 NA oil (WD=0.13mm) and filter set compatible with red (excitation wave length 555 nm, emission 630 nm). After imaging, the fluorescence signal was false colored in red using the NIS-element software to highlight bacterial membranes.

*Bacterial motility phenotypes*. The *P. fluorescens* SBW25 derivatives containing pPFL-dCas9 and pPFL-gRNA derivatives were grown overnight in LB medium containing kanamycin (50μg/ml) and gentamycin (10 μg/ml) at 28°C with shaking at 220 rpm. Overnight cultures from all strains were adjusted to OD_600_= 1 and a 1.5μ L drop was deposited at the center of the swarming plate (60mm x 15mm) containing semi-solid LB-agar (0.5%) supplemented with antibiotics and aT at 100ng/ml. Plates were incubated at 25°C for 48 hours before imaging. Swarming surface areas were assessed by imageJ (https://imagej.nih.gov/ij/). Surface areas of strains expressing gRNAs were normalized compared to the swarm area of the control strain containing a pPFL-gRNA empty control derivative with no guide RNA. Box plots of the distributions of numerical data obtained from ImageJ were displayed by Origin 8.3 software (https://www.originlab.com/).

*Survival after exposure to acute oxidative stress*. Strains were inoculated in LB supplemented with kanamycin 50μg/ml and gentamycin 10 μg/ml and grown overnight at 28°C with shaking at 220rpm. Cultures were then diluted to OD_600_ 0.02 in fresh media and allowed to grow up to OD_600_ 0.2-0.3 prior to addition of aT 100 ng/ml and additional cultivation for 4 hours at 28°C. All cultures were then adjusted to OD_600_=1 prior to addition of H_2_O_2_ at 2.5, 5 and10 mM final concentrations and further incubated for 30mn at 28°C. Cells were then collected, serially diluted in fresh LB and plated on selective LB agar plates. Bacterial viability was measured by counting the colony forming units after 24h at 28°C. Results were analyzed using Student’s t-test. P-value < 0.05 were considered statistically significant.

#### Biofilm formation

Cells carrying pPFL-dCa9 and pPFL-gRNA derivatives were grown at 28 °C in LB supplemented with kanamycin and gentamycin. Overnight cultures were diluted in fresh M9-glucose supplemented with the same antibiotics to OD_600_ 0.1 and grown at 28 °C for 5 hours. Cultures were adjusted to identical OD_600_ and inoculated to OD_600_=0.1 in triplicates in 12 wells culture plates containing fresh selective M9-glucose media supplemented with aT 100 ng/ml. Two additional wells were inoculated in the presence of Congo Red (CR). Culture plates were incubated at 25°C and 70% humidity in a growth chamber for 48h. Biofilm pellicles were carefully brought up to the top of the wells by slowly adding M9 media along the side of the well and then peeled off on a 25 mm diameter cover glass slide. The cover slides with intact biofilm pellicles were then mounted on an Attofluor™ Cell Chamber, covered with 1ml of PBS or transparent minimal media and stained with the FilmTracer FM 1-43 Green Biofilm dye (Molecular Probe) for 30 min prior to observation by confocal microscopy. Pellicles dyed with Congo red were mounted similarly and observed directly.

Stained biofilms were observed using a spinning disk confocal microscope (Nikon Eclipse Ti-E coupled with CREST X-Light^TM^ confocal imager; objectives Nikon CFI Plan Fluor 10X, DIC, 10x/0.3 NA (WD = 16 mm)). Excitation was performed at 470 nm and emission recorded at 505 nm (green). Congo Red stained cells were observed using excitation and emission wave lengths of 555 and 600 nm, respectively. Images were processed using IMARIS software (Bitplane, South Windsor, CT, United States). Biofilm images were quantified using the surface function in IMARIS (XTension biofilm). Means and maxima for surface thickness, roughness and surface substratum were determined from at least 3 independent pellicles and 2 measurements per pellicle Box plots of the distributions of numerical data obtained from Imaris were displayed using Origin 8.3 software.

## Supporting information

## Acknowledgements

This work was supported in part by funding through the Biological Systems Science Division, Office of Biological and Environmental Research, Office of Science, United States Department of Energy, under Contract DE-AC02-06CH11357.

## Author contributions statement

MFNG and P.N conceived the experiments; MFNG, S. F and G. M conducted the experiments; MFNG, SF, PEL and P. N analyzed the results. MFNG wrote the manuscript. All authors reviewed the manuscript.

