## Supplementary material for "Interrogation of genes controlling biofilm formation using CRISPR interference in *Pseudomonas fluorescens*"

Supplementary Information

**Supplementary Figure S1: CRISPR Interference System in *P. fluorescens*.**

**Supplementary Figure S2:** Determining optimal conditions for CRISPRi silencing in various *P. fluorescens* strains.

**Supplementary Figure S3:** Genomic organization of the genes targeted for silencing and annotation of protein functional domains**.**

**Supplementary Figure S4.** CRISPRi silencing of *ftsZ* and *mreB* using non-template strand gRNA.

**Supplementary Figure S5.** CRISPRi silencing of *ftsZ* and *mreB* using template strand gRNA.

**Supplementary Figure S6.** Effects of gRNA strand selection on swarming phenotypes.

**Supplementary Figure S7.** CRISPRi silencing of *gacS* increases cell survival to acute oxidative stress.

**Supplementary Table S1.** List of gBlocks gene fragments for gRNA designs.

**Supplementary Table S2.** Quantification of CRISPRi-mediated mNG gene silencing in *P. fluorescens*.

A

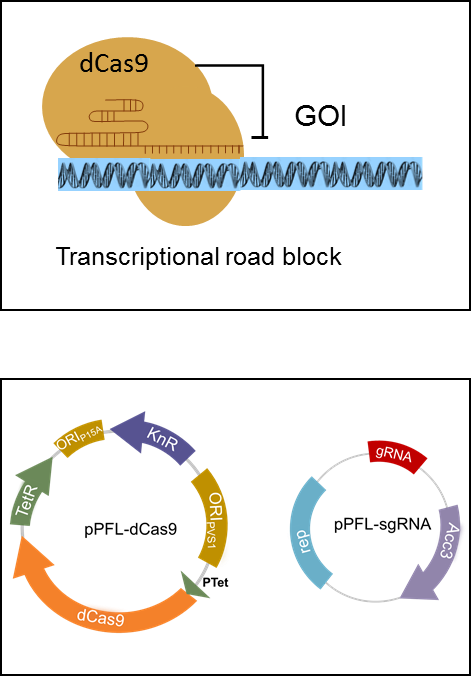

B

**Figure S1: CRISPR Interference System in *P. fluorescens***

**(A)** The catalytically inactive dCas9 protein binds the sgRNA chimera that will guide the protein-RNA complex to bind to a targeted gene and act as a transcriptional road block.

(B) The plasmid maps of the sgRNA and dCas9 expression vectors. The pPGL-sgRNA plasmid express the sgRNA from a constitutive promoter (pJ23119) with a strong terminator, rrnB, to ensure efficient termination, a gentamycin-selectable marker (Acc3) and a ColE1 replication origin, which is active in *E. coli* and *P. fluorescens*. The pPFL-dCas9 plasmid contains an aT-inducible promoter pLtetO-1 under control of the TetR repressor, a strong ribosomal binding site (RBS), a kanamycin-resistance marker (KnR), and two ORI regions composed of the P15A replication origin, functional in *E. coli* and the pVS1 replication origin functional in *P. fluorescens*.

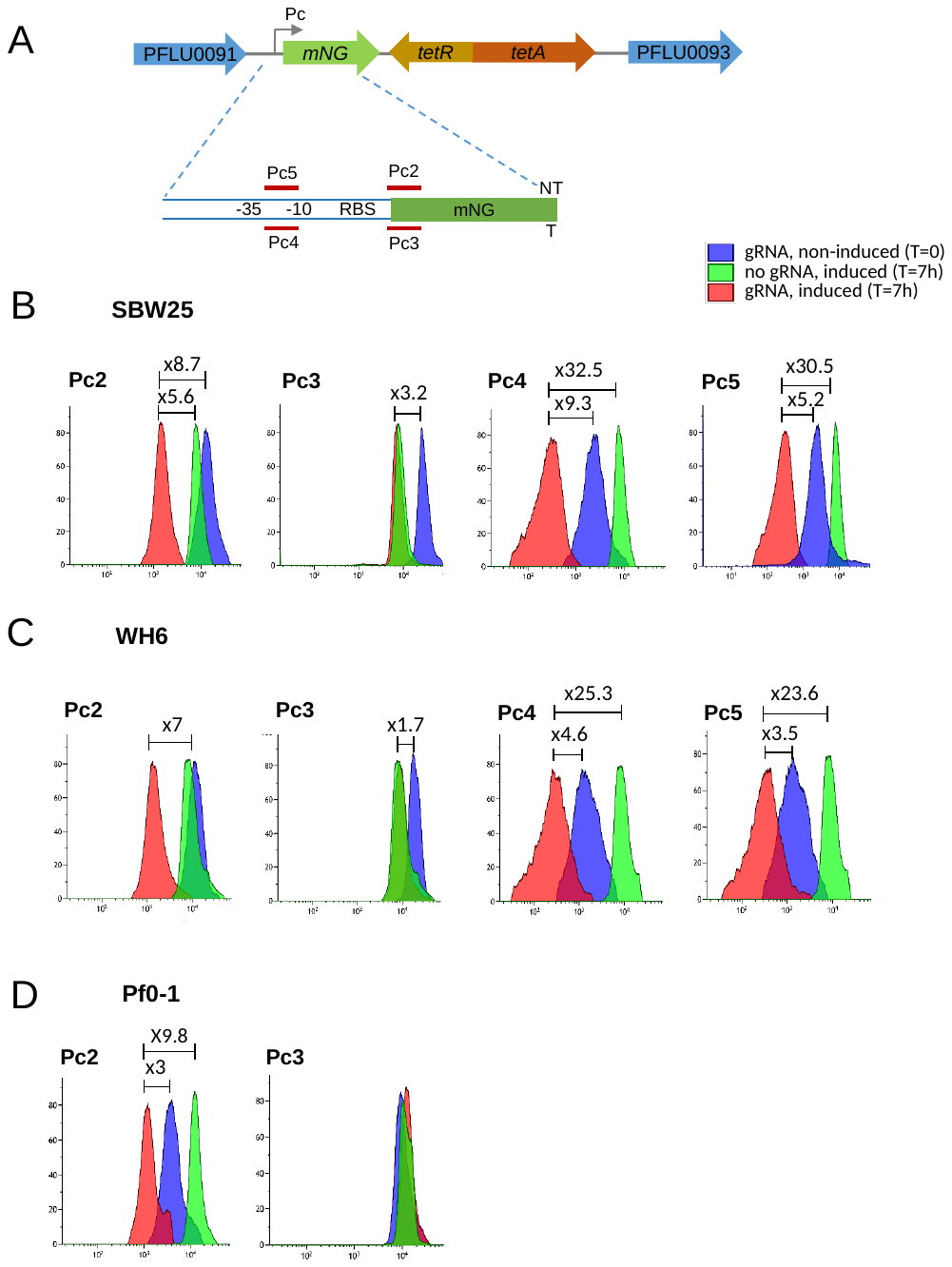

**Figure S2**:

**Figure S2**: **Determining optimal conditions for CRISPRi silencing in various P. *fluorescens* strains**

The *mNG* gene encoding the mNeonGreen fluorescent protein has been inserted at similar, presumably neutral, locations in the genomes of SBW25, WH6 and Pf0-1, as previously described ^1^. A) Genetic organization of the SBW25 chromosome surrounding the *Pc-mNG* construct used as a reporter for the activity of our CRISPRi system. A zoom-in section of the Pc region displays the positions targeted by the gRNAs (Pc2-5) relative to the promoter -35 and -10 motifs and the strand they copy. Expression of dCas9 was induced by addition of aT to the culture medium (T=0) and cells were analyzed by flow cytometry (CytoFlex S, Beckman) over time: (B) SBW25; (C) WH6; and (D) Pf0-1. At least 10^4^ particles were measured for each sample. Computerized gating in forward scatter (FSC) and side scatter (SSC) was used to eliminate cell debris. Histograms were generated using the Kaluza 2.0 software. In Pf0-1, peaks became very broad and overlapping when Pc4 and Pc5 gRNAs were used (data not shown). Cells expressing the Pc2 gRNA (NT strand, elongation block) displayed a decrease in fluorescence of 8-, 7- and 10-fold after 7 hours in SBW25, WH6 and Pf0-1, respectively, compared to cells expressing no guide. The silencing of *mNG* expression was much weaker (SBW25, WH6) or not detectable (Pf0-1) in cells expressing Pc3 gRNA (T strand, elongation block). When cells express the Pc4 and Pc5 gRNAs (initiation block), irrespective of T or NT strand, a large decrease in fluorescence was observed in SBW25 and WH6. Thus, repression by initiation-blocking guides in *P. fluorescens* appears to be independent of the targeted DNA strand, in keeping with previous observations in *E. coli*. Importantly, comparison of fluorescence intensities between no gRNA, induced (T=7h) and gRNA, non-induced (T=0) conditions reveals a lower fluorescence when gRNA is present in absence of inducer. This holds true for SBW25 and WH6 strains expressing Pc4 and Pc5, and for Pf0-1 expressing Pc2 gRNA. These findings suggest that dCas9 is expressed as a basal level in our system. Based on these observations, we selected Pc2-like gRNAs (NT strand, elongation block) for our experiments, unless otherwise indicated. Using Pc3-like gRNAs (T strand, elongation block) may provide a way for a moderate down-regulation of gene expression in SBW25 and WH6.

1- Noirot-Gros, M. F. *et al.* Dynamics of Aspen Roots Colonization by Pseudomonads Reveals Strain-Specific and Mycorrhizal-Specific Patterns of Biofilm Formation. *Frontiers in microbiology* **9**, 853, doi:10.3389/fmicb.2018.00853 (2018).

**Figure S3: Genomic organization of the genes targeted for silencing and annotation of protein functional domains.**

The organization of the genome around selected genes was extracted from the *P. fluorescens* SBW25 genome in Pseudomonas Genome DB (http://www.pseudomonas.com/). The positions of gRNAs copying the NT strand of targeted genes are indicated by red arrows. Protein Domain Architectures are from SMART (<http://smart.embl-heidelberg.de/smart>).

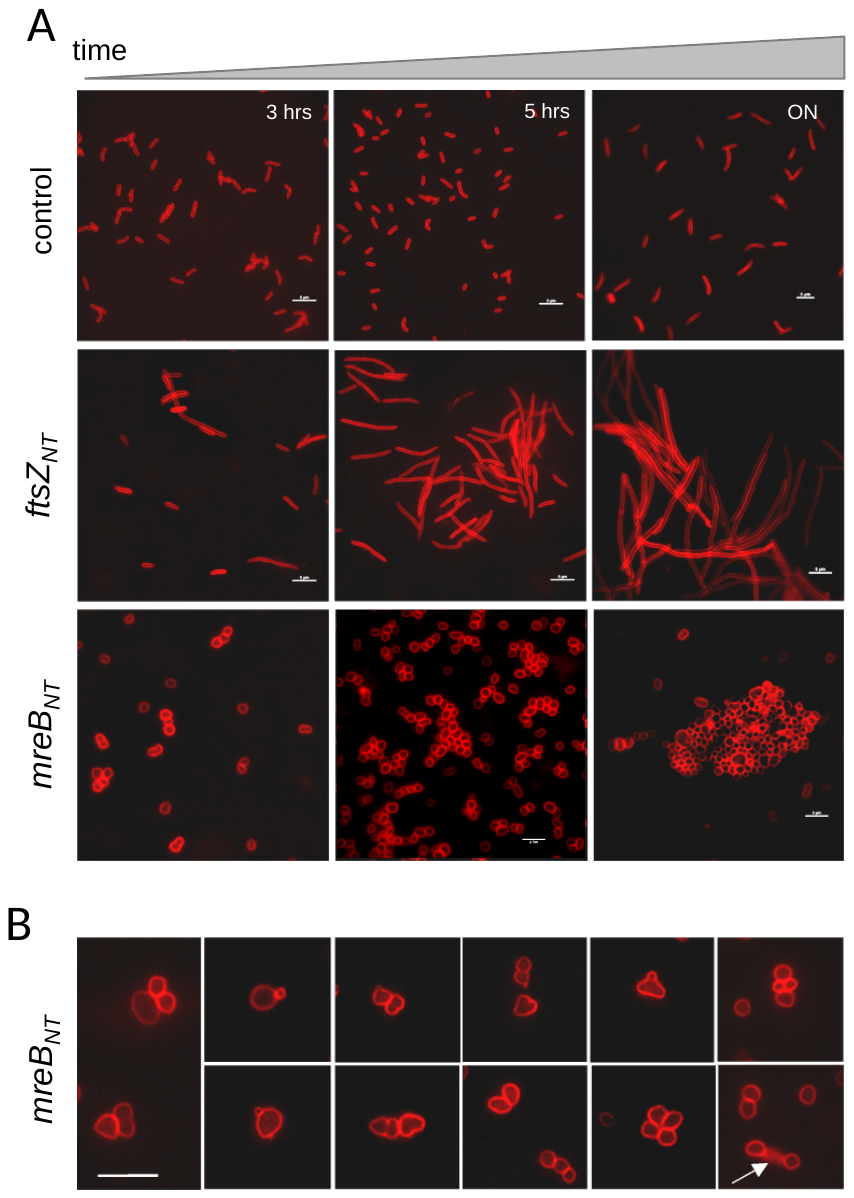

**Figure S4: CRISPRi silencing of *ftsZ* and *mreB* genes using non-template strand gRNA.**

(A) Cells carry pPFL-dCas9 and pPFL-*ftsZ*_NT_ (gRNA targeting *ftsZ* gene at non template strand) , pPFL-*mreB*_NT_ (gRNA targeting *mreB* gene at non template strand) or pPFL-control (no gRNA). dCas9 protein was induced by addition of 0.1μg/ml aT (final concentration). The morphological phenotypes were monitored by epifluorescence microscopy 3 hrs (left panels), 5 hrs (middle panels), and ON growth (18 hrs, right panels) after induction. Bacterial membranes were stained by the red fluorescent dye FM4-64 that incorporates into cell membranes. Scale bars correspond to 5μm. (B) Gallery of cells exhibiting morphological defects upon down regulation of *mreB*. White arrow points to a cell burst event.

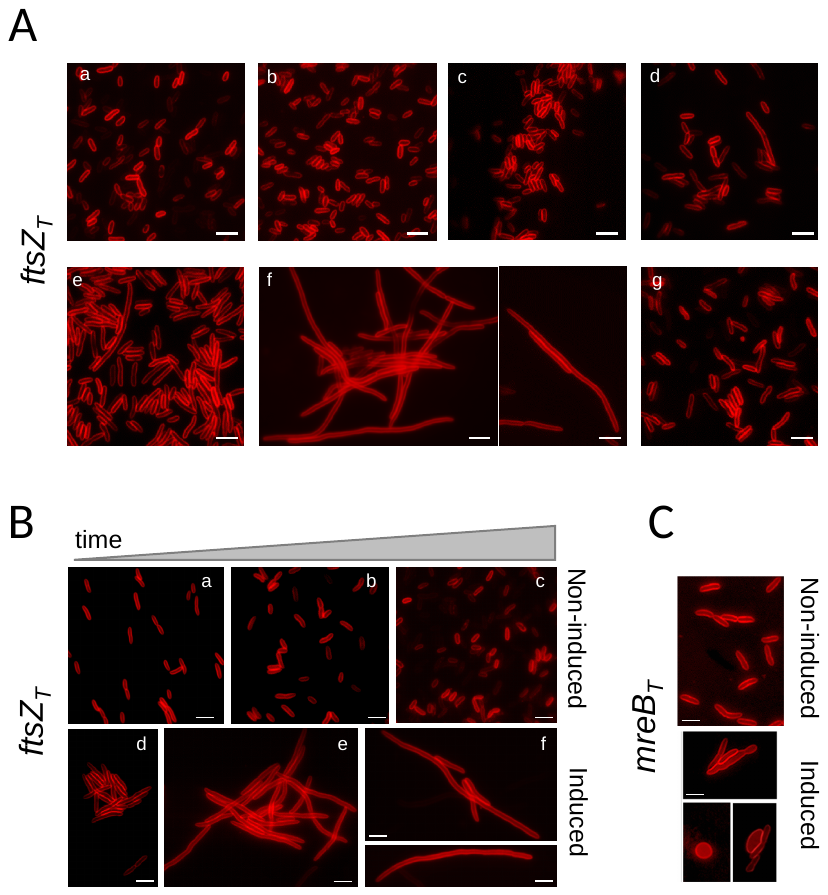

**Figure S5. CRISPRi silencing of *ftsZ* and *mreB* genes using template strand gRNA.**

Dose responses were monitored in cells carrying pPFL-dCas9 and pPFL-*ftsZ*_T_ (gRNA targeting *ftsZ* gene at template strand) the pPFL-control (no gRNA). dCas9 was induced with increasing amounts of aT: (a), 0; (b), 0.02; (c), 0.05; (d), 0.1; (e), 0.2; (f), 0.5 μg/ml; and (g) control strain treated with 0.5 μg/ml aT. The filamentation phenotype was monitored by epifluorescence microscopy 7 hours after induction. Bacterial membranes were stained with FM4-64. Scale bars correspond to 2μm. (**B**) Induction of the filamentation phenotype over time. Cells expressing the *ftsZ_T_* guide were either not induced (a, b and c) or induced with aT 0.5μg/ml (d, e and f) for dCas9 expression. Cells were stained with FM4-64 and observed by epifluorescence microscopy after 3 hours (a, d), 7 hours (b, e) and overnight (c, f) post induction. Scale bars correspond to 2μm. (**C**) Cells expressing the *mreB*_T_ guide (targeting *mreB* gene at template strand) were induced with 0.5 μg/ml aT and morphological phenotypes were observed after ON growth (18 hrs).

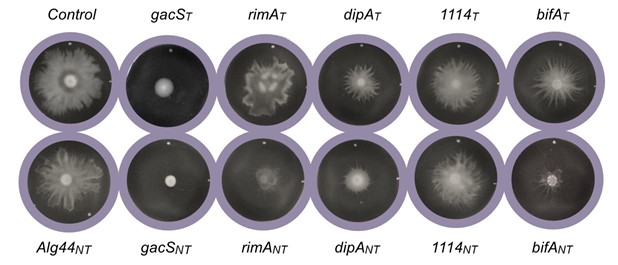

**Figure S6. Effects of gRNA strand selection on swarming phenotypes.**

*P. fluorescens* strains harboring pFL-dCas9 and various pFL-gRNA expressing either gRNA_NT_ or gRNA_T_ for each targeted gene. In experiments involving gRNA_NT_ and gRNA_T_, dCas9 was induced by 0.1μg/ml and 0.5 μg/ml aT (final concentrations), respectively. gRNAs target the start of the coding regions of *gacS* (PFLU3777) as well as of genes encoding c-di-GMP binding proteins, *rimA* (PFLU0263), *dipA* (PFLU0458), *alg44* (PFLU0988), *bifA* (PFLU4858) and PFLU1114 (see also Fig. 6). Control corresponds to the pFL-gRNA plasmid with no guide RNA inserted.

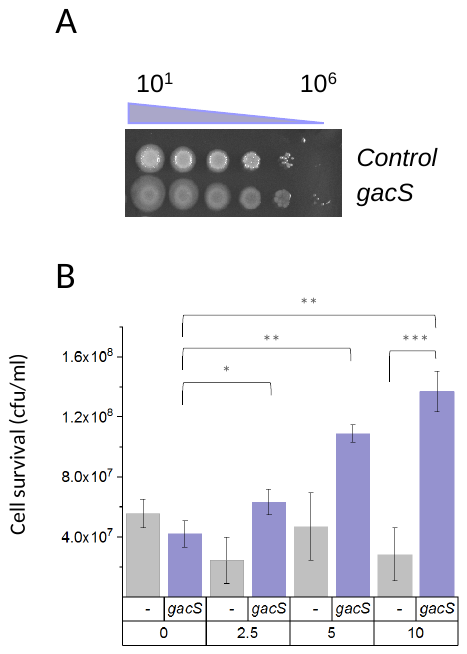

**Figure S7: CRISPRi silencing of *gacS* increases cell survival to acute oxidative stress**

*P. fluorescens* strain harboring pPFL-dCas9 and expressing the *gacS_NT_* guide to downregulate the two-component kinase gene *gacS* (PFLU3777). Control (-) corresponds to the pFL-gRNA plasmid with no gRNA inserted. Expression of dCas9 was induced 4h prior to exposure of cells to various H_2_O_2_ concentrations for 30 min. Treated cultures were then serially diluted in LB. (A) Dilutions were either spotted onto LB agar plates containing Kn and Gm and incubated for 24 hrs at 28°C. (B) Dilutions were plated on the same medium for colony forming unit (cfu) counts. Cell survival is the number of cfu/ml after 24hrs at 28°C. Experiments were repeated three times independently for each strain and each H_2_O_2_ treatment. Error bars are standard deviation of the means. * P> 0.05, ** P >0.005, *** P>0.0005.

| gene ID / NCBI | Target gene | Strand | Target sequence |
| --- | --- | --- | --- |
| Pc-mNG | Pc2 | NT | TTCATGAGTTGCCGGTAAAC |
| Pc-mNG | Pc3 | T | GATATACATATGAATTCGAA |
| Pc-mNG | Pc4 | T | CGGTCTGTAGGCTGTAATGC |
| Pc-mNG | Pc5 | NT | TTACAGCCTACAGACCGAGA |
| PFLU_RS04730 | pflu_0952 | T | AGACAACATCCCCGCCAGCC |
| PFLU_RS04731 | pflu_0952 | NT | CACTTTGATGACCGGGCTGG |
| PFLU_RS04305 | pflu_0863 | T | CAGCGATCTTTCCATTGACC |
| PFLU_RS04306 | pflu_0863 | NT | CAGGTCAATGGAAAGATCGC |
| PFLU_RS18410 | pflu_3777 | T | CTGACAAGAATGGGGATAAA |
| PFLU_RS18410 | pflu_3777 | NT | ATAACGTCAGCAACAGTACG |
| PFLU_RS01300 | pflu_0263 | T | CTCGCCCAACCAGCGCTGCG |
| PFLU_RS01300 | pflu_0263 | NT | GGTGAGTGAAAGGGGGAAGT |
| PFLU_RS02265 | pflu_0458 | T | CCAGCCCGATGTCGCCCGAA |
| PFLU_RS02265 | pflu_0458 | NT | GCCGCCATTCGGGCGACATC |
| PFLU_RS04905 | pflu_0988 | NT | TGGACTACGTTGGCATTTAC |
| PFLU_RS23815 | pflu_4858 | T | GGAACTCAAGAACAGCTTGT |
| PFLU_RS23815 | pflu_4858 | NT | GCCGCCATTCGGGCGACATC |
|  | pflu_1114 | T | TATGACCCCGGATTCAGTCA |
|  | pflu_1114 | NT | TGACTGAATCCGGGGTCATA |
| Block sequence backbone | GCTCAGTCCTAGGTATAATACTAGTNNNNNNNNNNNNNNNNNNNNGTTTTAGAGCTAGAAATAGCAAGTTAAAATAAGGCTAGTCCGTTATCAACTTGAAAAAGTGGCACCGAGTCGGTGCTTTTTTTGAAGCTTGGGCCCGAAC | | |

**Table S1: List of gBlocks gene fragments for gRNA designs.**

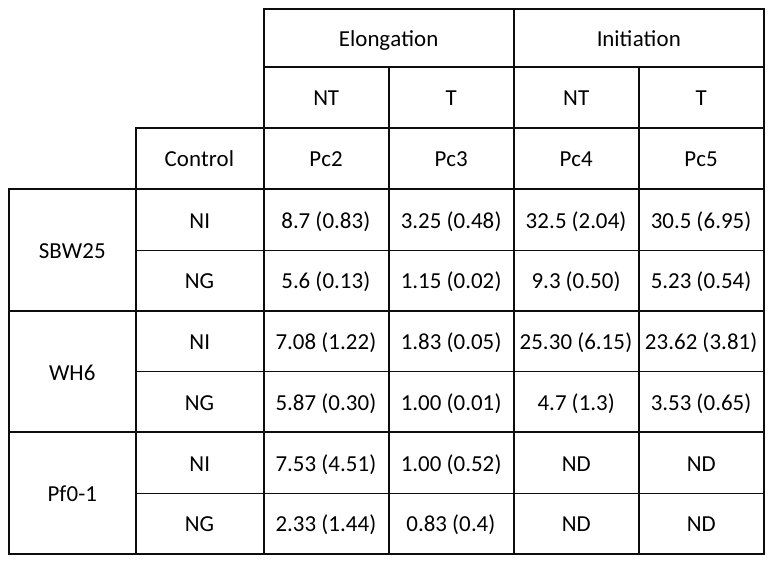

**Table S2: Quantification of mNG gene silencing by CRISPRi in *P. fluorescens*.**

Fluorescence in cultures of SBW25, WH6 and Pf0-1 cells expressing *Pc-mNG* was measured as the median of the flow cytometry peaks 7 hours after induction of dCas9 by addition of 0.1µg/ml aT (see Figure S2). Gene silencing can be measured relative to a control which is the same strain without dCas9 induction (non-induced, NI) or relative to a control strain without gRNA but in which dCas9 is induced (no-guide, NG). gRNAs were copied from the non-template (NT) or template (T) DNA strand and targeted transcription initiation or elongation block (Figure S2). Mean values of fluorescence decrease were measured from independent experiments (n ≥ 3) and standard deviation of the means are in parenthesis. ND, not determined.
